# π-SPECFormer: A Mass Spectrometry-based Foundation Model for Decoding the Dark Proteome

**DOI:** 10.64898/2026.11.06.764812

**Authors:** Tianze Ling, Tao Liu, Fan Xu, Yiming Zhang, Jun Xia, Xiansong Huang, Shaorong Chen, Linhai Xie, Rong Xu, Tingpeng Yang, Kaixuan Li, Guibin Wang, Zhendong Liang, Jun Zhu, Stan Z. Li, Fuchu He, Yu Wang, Cheng Chang

## Abstract

In mass-spectrometry-based proteomics, a substantial proportion of acquired spectra remain unidentified, concealing a vast reservoir of biological variations commonly termed the “dark proteome”, potentially obscuring sequence mutations, non-canonical peptides and post-translational modifications. Existing deep-learning methods largely rely on supervised training with peptide–spectrum matches (PSMs), limiting their ability to learn from the much larger pool of unannotated spectra. Here we introduce π-SPECFormer, a mass-spectrometry foundation model pre-trained on 291 million spectra through a two-stage framework that combines self-supervised and PSMs-guided representation learning. π-SPECFormer transfers effectively to *de novo* peptide sequencing, spectrum clustering and modification detection, consistently outperforming task-specific baselines. When applied to proteomic profiles from 103 individuals with lung adenocarcinoma, the model identified nine high-confidence mutated peptides, including six that were absent from the reference databases. π-SPECFormer thus enables unbiased decoding of molecular complexity beyond the boundaries of the canonical proteome.

## Main

In shotgun proteomics, proteins are enzymatically digested into peptides and analyzed by liquid chromatography coupled with tandem mass spectrometry (LC-MS/MS)^1–3^. Among existing peptide identification strategies, database searching remains the dominant approach. This strategy identifies peptides by comparing experimentally acquired tandem mass spectra with theoretical fragment ions predicted from a reference protein sequence database^4^. This approach is constrained by the completeness of the sequence database and by the range of modifications included in the search. Consequently, spectra derived from sequence mutations, non-canonical peptides or unexpected post-translational modifications (PTMs) can remain unassigned. Indeed, it has been reported that fewer than half of acquired spectra typically receive high-confidence peptide assignments (**Fig. 1a**)^5^. To further confirm this limitation, we analyzed the ten largest datasets in π-MSNet^6^—a large-scale, AI-ready proteomics data resource—and found that the spectral identification rates consistently ranged between ∼20% and 30% (**Fig. 1b**), underscoring the broad relevance of this challenge.

**Fig. 1.**
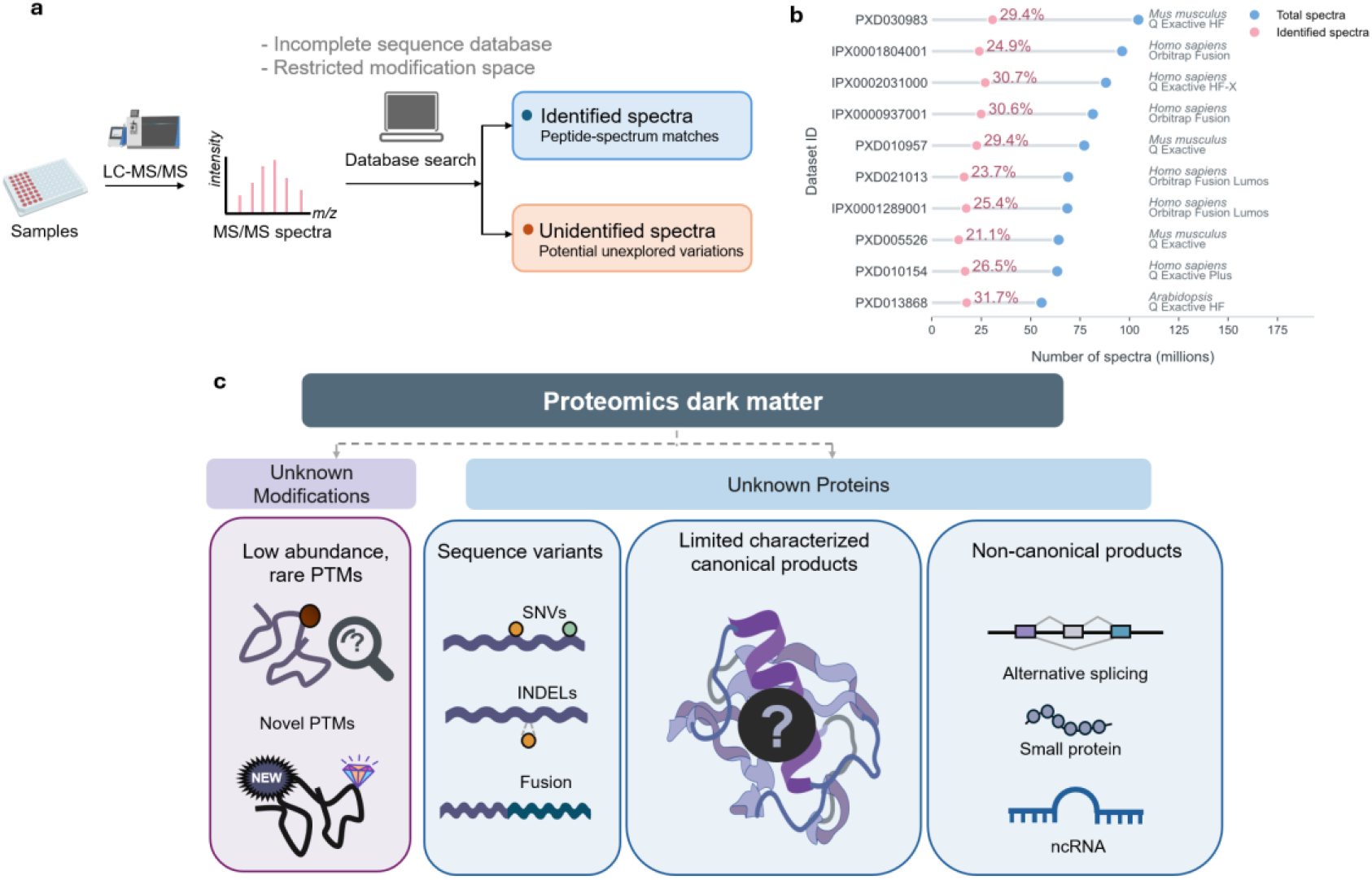
The landscape of proteomic dark matter: origins, prevalence, and classification. **a**, Origins of proteomic dark matter. Schematic of conventional database-based peptide identification in shotgun proteomics. Tandem mass spectra acquired by LC–MS/MS are matched to theoretical fragment ions generated from a reference sequence database. Incomplete sequence coverage and restricted modification search spaces leave many spectra unassigned (dark matter). Blue and orange indicate identified and unidentified spectra, respectively. **b,** Prevalence of dark matter across large-scale datasets. By re-analyzing the ten largest datasets included in π-MSNet using conventional approaches (as depicted in a), we observed consistently low identification rates. Blue and pink circles denote total and identified spectra, respectively, and connecting lines indicate the corresponding unassigned spectral space. Percentages indicate the proportion of spectra assigned to peptide sequences in each dataset (ranging from ∼20% to ∼31%). Dataset identifiers are shown on the left, with the corresponding species and mass-spectrometer type indicated on the right. **c,** Operational classification of proteomic dark matter. Dark matter is categorized into uncharacterized modifications and uncharacterized protein forms. The latter comprise poorly characterized canonical products and non-canonical products derived from mutations (e.g., SNVs and indels), alternative splicing and ncRNAs. SNV, single-nucleotide variant; indel, insertion or deletion; PTM, post-translational modification; ncRNA, non-coding RNA.

Unidentified spectra can contain biologically relevant molecular information. Deep proteome profiling of HeLa cells has revealed thousands of protein-level sequence mutations and alternative-splicing products, showing that up to 73% of non-synonymous mutations detected at the RNA level are indeed translated into protein^7^. Open searches with a mass tolerance window of ±500 Da had further shown that unexpected and substoichiometric modifications account for at least one-third of unidentified spectra missed by conventional searches^5^. Recently, proteogenomic studies have identified 8,945 previously unannotated microproteins encoded by small open reading frames (smORFs) from gastric cancer tissues, which may influence the regulation of cancer-cell proliferation^8^. Meanwhile, the TransCODE Consortium generated a consensus landscape of protein-level evidence for non-canonical open reading frames (ncORFs) by analyzing 95,520 proteomics experiments, finding that about 25% of 7,264 ncORFs yield detectable peptides, defining “peptideins” to advance biomedical discovery across understudied components of the human proteome^9^. Together, these observations indicate that unassigned spectra constitute not only technical or analytical failures, but also a potential source of previously inaccessible proteomic variation.

We term this body of uncharacterized proteomic data the “dark matter” of the proteome—an analogy borrowed from astronomy. In astronomy, the Swiss astronomer Fritz Zwicky inferred the existence of dark matter from the unexpectedly high velocities of galaxies in clusters, which could not be explained by visible mass alone^10^. In recent years, different forms of uncharacterized proteomic information have collectively been described as proteomic “dark matter”. Previous definitions have emphasized unexpected modifications^5^, proteins lacking structural characterization^11^ or translation products derived from non-canonical open reading frames and nominally non-coding RNAs^12^. Here, we use proteomic “dark matter” as a systematic and unified definition term for molecular entities that are present in biological samples but remain difficult to identify or characterize using current experimental and computational workflows or whose biological functions remain uncharacterized (**Fig. 1c**). We distinguish two broad classes: uncharacterized protein forms and uncharacterized modifications. The former includes mutation peptides arising from genomic alterations, poorly characterized products of canonical genes, and products of non-canonical transcription or translation.

Deep learning has enabled increasingly powerful analysis of large-scale mass-spectrometry data by learning informative representations from high-dimensional spectra^3, 13, 14^. Existing approaches in this domain broadly fall into two categories: “peptide-to-spectrum”^15–19^ and “spectrum-to-peptide”^20–27^ models, demonstrating strong model performance. However, most existing models are trained for individual tasks using PSM data (which is “labeled data”) generated by conventional identification pipelines. These labels are restricted to the sequence and modification spaces encoded by the corresponding searches and can vary across search engines and parameter settings^28–30^. Large-scale representation learning directly from unannotated spectra offers an alternative: it can capture general fragmentation patterns without requiring peptide identities and may yield transferable representations for interrogating proteomic features beyond predefined search spaces.

These limitations motivate us to develop a deep learning model specifically designed for proteomic dark matter, capable of directly learning the signal distributions embedded within unannotated spectra. To the best of our knowledge, although pre-trained large models have found broad application across the life sciences—including protein language modeling, protein design, metabolite discovery and single-cell omics analysis^31–35^. To our knowledge, few studies have explored fully self-supervised pre-training directly on proteomics tandem mass spectra at this scale^36^. Here we introduce π-SPECFormer, a mass-spectrometry foundation model pre-trained on 291 million tandem mass spectra using a two-stage strategy that combines self-supervised learning with PSM-guided representation learning. π-SPECFormer learns transferable spectrum embeddings that can be adapted to diverse analytical tasks. Across benchmarks for *de novo* peptide sequencing, spectrum clustering and PTMs detection, the pre-trained model consistently outperformed task-specific baselines. We further applied π-SPECFormer to proteomic profiles from 103 individuals with lung adenocarcinoma, identifying nine high-confidence mutation peptides, including six mutations absent from the public databases examined. Synthetic-peptide validations supported their spectral assignments, and integrative clinical and structural analyses linked these mutations to patient outcomes or predicted alterations in protein interactions. Together, these results establish π-SPECFormer as a generalizable framework for spectrum representation learning and for extending proteomic analysis beyond conventional database searches.

## Results

### Two-stage contrastive pre-training learns transferable spectrum representations

The π-SPECFormer pre-training framework comprises two consecutive stages (**Fig. 2a**). In the first stage, we generated two independently augmented views of each unlabeled tandem mass spectrum by randomly masking signal peaks. One view was processed by an online spectrum encoder to produce a query embedding, whereas the other was processed by a momentum-updated encoder to produce the corresponding target embedding. Embeddings of previously processed spectra were retained in a dynamic queue and used as negative keys. Both encoders comprised a peak-embedding module followed by a Transformer backbone, but gradients were propagated only through the online encoder; the momentum encoder was updated as an exponential moving average of the online parameters. We optimized an InfoNCE objective that increased the similarity between embeddings of two masked views derived from the same spectrum relative to embeddings of other spectra in the queue. This design encouraged the model to learn spectrum-level representations that were robust to partial peak removal.

**Fig. 2.**
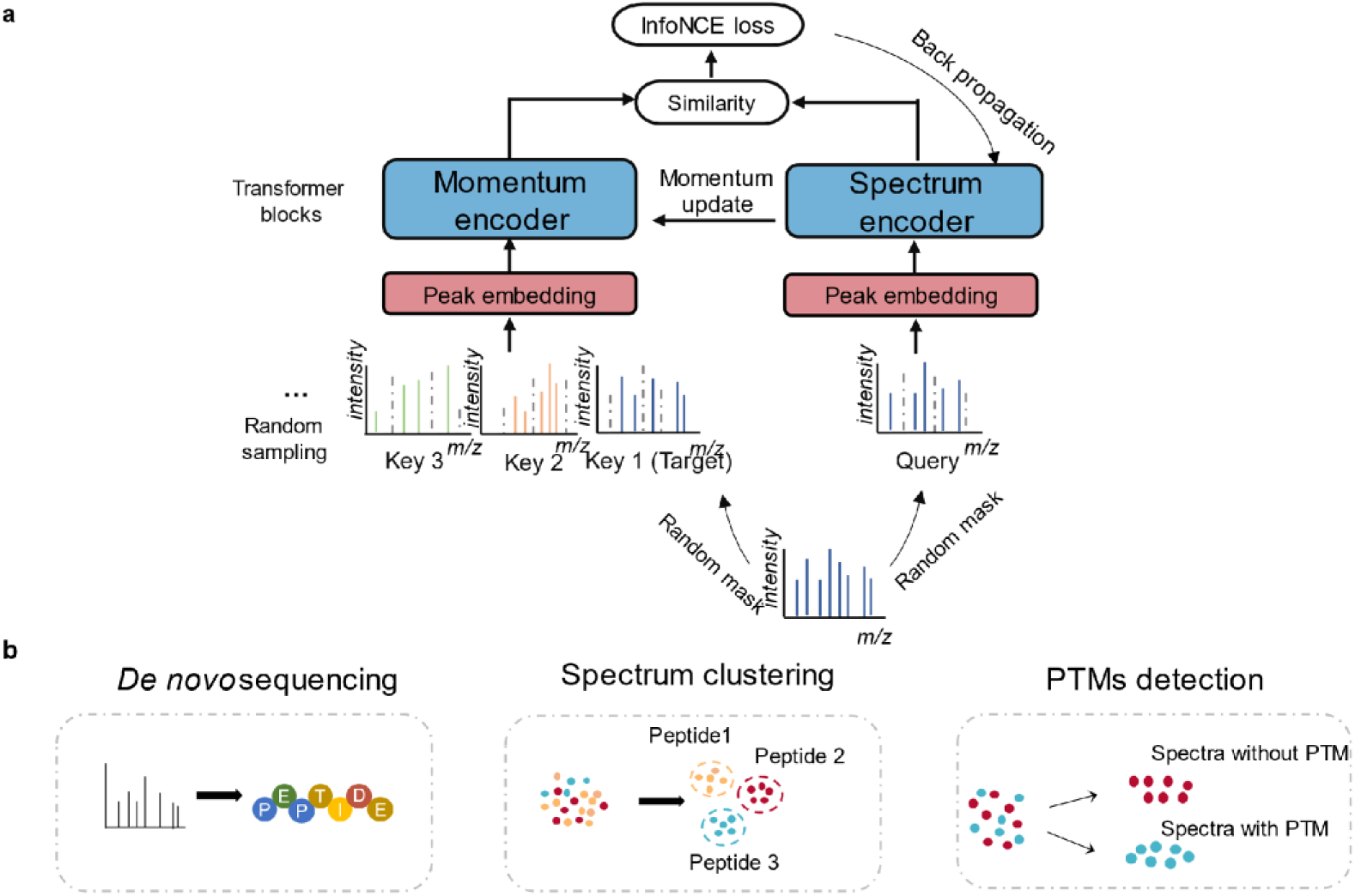
Schematic overview of the π-SPECFormer architecture. **a**, Self-supervised pre-training phase. Unlabeled mass spectra underwent a dual-masking strategy to generate augmented spectra. Within a contrastive learning framework, the model learned robust spectral representations by matching an encoded query spectrum (Query) to a dictionary of encoded key spectra (Key1, Key2, Key3…) using the InfoNCE (Information Noise Contrastive Estimation) loss^37^. **b,** Deployment of the pre-trained spectrum encoder in downstream applications. The learned representations are transferred to de novo peptide sequencing, spectrum clustering and PTMs detection using task-specific decoding, clustering and classification procedures, respectively.

In the second stage, we refined the pre-trained representation using PSM-informed spectrum grouping. Spectra assigned to the same precursor were treated as positive pairs, whereas spectra with distinct assignments were used as negative examples. Peptide identities were used to construct spectrum pairs but were not provided as model inputs. The model architecture and contrastive objective were otherwise unchanged. We refer to the checkpoints obtained after the self-supervised and PSM-guided stages as π-SPECFormer-Stage 1 and π-SPECFormer-Stage 2, respectively. Full details of spectrum preprocessing, pair construction and momentum updates are provided in **Methods**.

To track the downstream utility of the learned representations during pre-training, we evaluated phosphorylation detection at pre-defined training checkpoints using a fixed data split and an identical evaluation protocol. Phosphorylation-detection accuracy (determining whether phosphorylation is present in the mass spectrum, IPX0001804002 dataset) increased as the contrastive loss decreased and plateaued at later checkpoints in both pre-training stages. This concordant trend indicates that contrastive pre-training progressively enriched the spectrum embeddings with features informative for modification detection. Importantly, phosphorylation labels were not used to optimize the spectrum encoders during pre-training.

We also evaluated masked-peak reconstruction as an alternative self-supervised objective. In this setting, selected peak attributes were masked and predicted from the remaining spectrum. Although the reconstruction loss rapidly reached a plateau, this pre-training strategy did not improve performance in the downstream tasks examined.

We next adapted the pre-trained spectrum encoder to three downstream applications: *de novo* peptide sequencing, spectrum clustering and PTMs detection (**Fig. 2b**). The same pre-trained backbone provided spectrum embeddings for each application, together with task-specific decoding, clustering or classification procedures.

### Pre-trained spectrum representations support diverse downstream proteomics tasks

We evaluated the transferability of π-SPECFormer representations across three downstream tasks: *de novo* peptide sequencing, spectrum clustering and PTMs detection (**Fig. 3a, b**). These tasks require complementary aspects of spectrum representation learning. *De novo* sequencing requires the recovery of sequence-informative fragmentation patterns, spectrum clustering assesses whether embeddings capture peptide-level spectral similarity, and PTMs detection requires the recognition of modification-associated fragmentation features. Together, they provide a broad evaluation of whether the representations learned during pretraining can support distinct spectrum-analysis tasks.

**Fig. 3.**
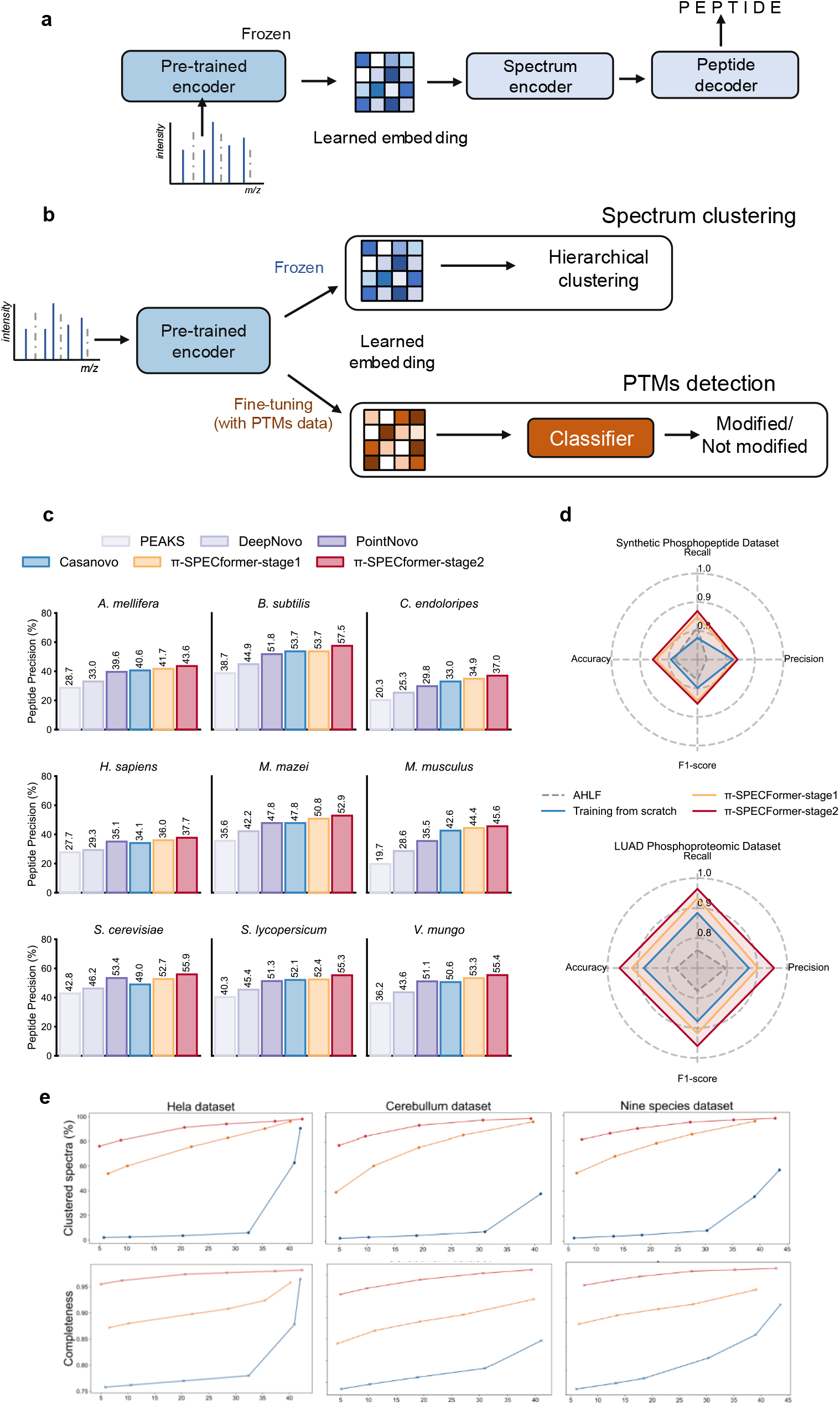
Pre-trained π-SPECFormer representations improve diverse downstream proteomics tasks. **a**, Adaptation of the pre-trained π-SPECFormer encoder to *de novo* peptide sequencing. The parameters of the pre-trained model are frozen, and its peak-level representations are supplied to a task-specific spectrum encoder and autoregressive peptide decoder to generate complete peptide sequences. **b,** Adaptation of π-SPECFormer to spectrum clustering and PTMs detection. For spectrum clustering, embeddings generated by the frozen pre-trained model are subjected to hierarchical clustering. For PTMs detection, the pre-trained model is fine-tuned using PTM-labelled spectra together with a classification head to distinguish modified from unmodified spectra. **c,** Peptide precision of PEAKS, DeepNovo, PointNovo, Casanovo, π-SPECFormer-Stage 1 and π-SPECFormer-Stage 2 on the nine-species *de novo* sequencing benchmark. A peptide prediction is considered correct only when the complete predicted sequence matches the reference sequence. Values above the bars indicate peptide precision. **d,** Performance of AHLF, a model trained from scratch, π-SPECFormer-Stage 1 and π-SPECFormer-Stage 2 for phosphorylation-spectrum detection on the synthetic phosphopeptide dataset and the LUAD phosphoproteomic dataset. Radar plots show recall, precision, F1 score and accuracy. **e,** Spectrum-clustering performance on the HeLa, cerebellum and nine-species datasets. The upper panels show the proportion of spectra assigned to non-singleton clusters, and the lower panels show cluster completeness, both as functions of the proportion of incorrectly clustered spectra. Results are shown for GLEAMS, π-SPECFormer-Stage 1 and π-SPECFormer-Stage 2.

*De novo* peptide sequencing directly infers peptide sequences from tandem mass spectra without relying on a protein database. It therefore provides a complementary strategy for detecting sequence mutations, modified peptides and peptides derived from genomic regions outside conventional protein annotations^38, 39^. To adapt π-SPECFormer to *de novo* sequencing, we used the pre-trained model as a frozen spectrum encoder (**Fig. 3a**), which is inspired by the work of Zhu et al.^40^ The parameters of the pre-trained model were fixed during downstream training, and its peak-level representations were supplied to a task-specific Transformer encoder and autoregressive peptide decoder. The resulting models are referred to as π-SPECFormer-Stage 1 and π-SPECFormer-Stage 2 according to the pretraining checkpoint used for feature extraction.

We first evaluated both models using the nine-species benchmark originally introduced with DeepNovo—the pioneering deep learning-based *de novo* sequencing model^19^. We compared π-SPECFormer with several representative commonly used *de novo* sequencing tools, including PEAKS^41^, DeepNovo^27^, PointNovo^21^, and Casanovo^25^ (**Fig. 3c**). Because a peptide-level prediction was considered correct only when the complete predicted sequence matched the reference sequence, we used peptide precision as the primary evaluation metric. π-SPECFormer-Stage 1 achieved higher peptide precision than all four reference methods in eight of the nine datasets and matched the best baseline in the remaining dataset. Relative to Casanovo, the strongest baseline overall, Stage 1 improved peptide precision by as much as 7.6%, corresponding to an increase from 49.0% to 52.7% in *Saccharomyces cerevisiae*. Stage 2 achieved the highest peptide precision in all nine datasets. Compared with the strongest non-π-SPECFormer baseline in each dataset, its largest relative improvement was 12.1%, from 33.0% to 37.0% in *C. endoloripes*. These results indicate that both self-supervised and PSMs-guided pre-training improved complete-sequence prediction.

We next examined whether the learned spectrum embeddings supported peptide-level spectrum clustering (**Fig. 3b**). Spectrum clustering groups tandem mass spectra according to their fragmentation patterns and can facilitate the propagation of peptide annotations to unassigned spectra, the detection of inconsistent assignments and the identification of spectral outliers^42^. Clustering performance was evaluated by varying the clustering threshold and measuring the proportion of spectra assigned to non-singleton clusters and cluster completeness as functions of the incorrectly clustered spectrum rate. Across the HeLa, cerebellum and nine-species datasets, both π-SPECFormer checkpoints produced more favorable clustering curves than GLEAMS^43^, recovering a larger proportion of spectra and higher cluster completeness at matched error rates (**Fig. 3e**). At a clustered-spectrum proportion of 50%, π-SPECFormer-Stage 2 reduced the incorrectly clustered spectrum rate by 99.4%, 99.5% and 99.5% relative to GLEAMS in the HeLa, cerebellum and nine-species datasets, respectively. Stage 2 also consistently outperformed Stage 1, indicating that PSM-guided refinement further improved the separation of spectra assigned to different peptides.

For PTMs detection, we initialized the spectrum encoder using the pre-trained π-SPECFormer checkpoints, attached a classification head and fine-tuned the resulting model using PTM-labelled spectra to distinguish modified from unmodified spectra (**Fig. 3b**). We compared π-SPECFormer with AHLF, a convolutional neural network-based phosphorylation spectrum detection model, currently one of the top-performing models for this task^44^ using precision, recall, F1 score and accuracy. On the synthetic phosphopeptide dataset, π-SPECFormer-Stage 2 outperformed AHLF across all four metrics, increasing the F1 score and accuracy by 7.7% and 8.4%, respectively (**Fig. 3d**). The advantage was more pronounced on the LUAD phosphoproteomic dataset, on which Stage 2 increased the F1 score by 23.6% and accuracy by 24.3% relative to AHLF. Stage 2 also outperformed the same downstream architecture trained from scratch, demonstrating that large-scale spectrum pre-training improved modification detection beyond the contribution of the downstream architecture alone.

Together, these results show that π-SPECFormer learns transferable spectrum representations that support three complementary forms of spectrum analysis. Frozen peak-level representations improve complete peptide-sequence prediction, spectrum-level embeddings enable accurate peptide-level clustering without sequence information, and task-specific fine-tuning supports the detection of phosphorylation and diverse PTMs directly from tandem mass spectra. The consistent improvements across species, biological samples, proteases and modification classes indicate that large-scale pre-training captures transferable fragmentation features that are not restricted to a single experimental setting or downstream task.

### π-SPECFormer uncovers a novel functional and prognostic protein mutation in lung adenocarcinoma

We next evaluated whether π-SPECFormer could reveal biologically and clinically informative components of proteomic dark matter. We focused on discovering single-amino-acid variants in a clinical tumor cohort. This focus is motivated by the functional significance of mutations: some can alter key cellular functions, and the accumulation of such functionally consequential mutations is a central mechanism of cancer development^45^. We reanalyzed a dataset from paired tumor and adjacent non-tumor tissues collected from 103 individuals in the CNHPP-LUAD cohort^46^ (**Fig. 4a**). We developed a *de novo*-sequencing-based workflow that combined sequence prediction, reference-proteome alignment, peptide-spectrum quality control and quantitative analysis. This analysis identified 5,103 candidate single-amino-acid mutations across 1,712 proteins. We systematically annotated the candidate mutations using dbSNP^47^, GDC^48^, COSMIC^49^ and literature in PubMed^50^ to determine whether corresponding nucleotide or amino-acid changes had previously been reported as population polymorphisms or disease-associated mutations. Of the 5,103 candidates, 3,946 (77.3%) were not represented in the resources examined, 1,089 (21.3%) matched previously reported rare mutations with a minor allele frequency of no more than 1% in at least one population, and 68 (1.3%) corresponded to common polymorphisms with a minor allele frequency greater than 1% (**Fig. 4b**).

**Fig. 4.**
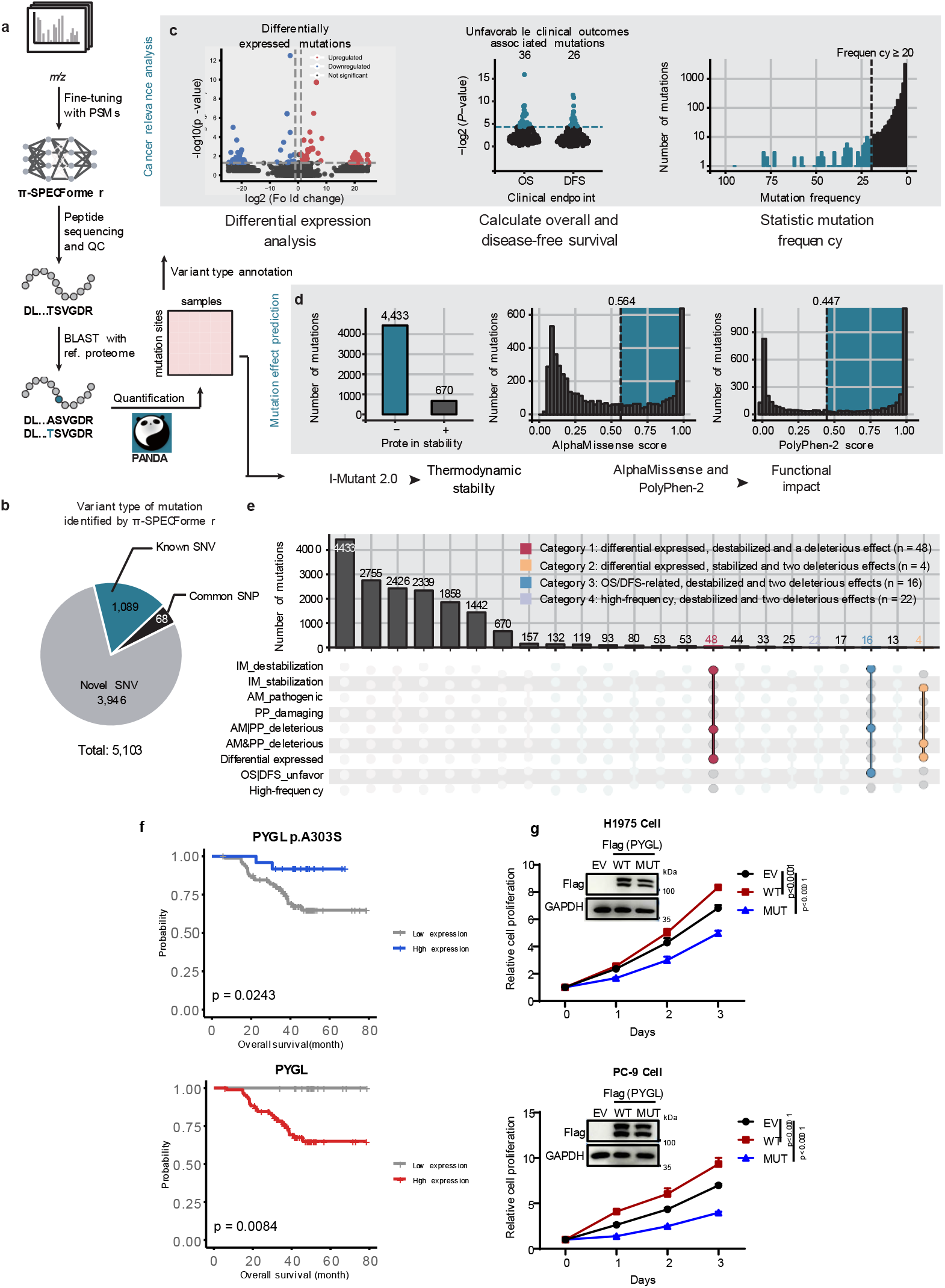
Identification of cancer-related mutations in 103 patients from the CNHPP-LUAD cohort using π-SPECFormer. **a**, Schematic overview of the π-SPECFormer workflow designed to uncover 5,103 substitution mutations unidentified in the previous research. **b,** Distribution of mutations according to genetic polymorphism status. **c,** Cancer relevance analysis, including differential expression of mutated proteins, clinical risk evaluation and mutation frequency profiling. Differential expression was assessed using paired t-tests with fold-change cutoffs of > 2 or < 0.5. Associations with overall survival (OS) and disease-free survival (DFS) were evaluated by Cox proportional hazards regression, with the optimal stratification cut-off determined by MaxStat and significance assessed by the log-rank test. For mutation frequency analysis, a mutation was scored as present in a patient if it was detected in either tumor or adjacent non-tumor tissue. **d,** Mutation effect prediction, including thermodynamic stability, AlphaMissense and PolyPhen-2. The cutoff of deleterious effect for AlphaMissense and PolyPhen-2 is 0.564 and 0.447 according to official user manual. **e,** Upset plots illustrate the inclusive intersections of four categories of mutations with multidimensional criteria. IM, I-Mutant2.0; AM, AlphaMissense; PP, PolyPhen-2. **f**, OS curve of PYGL p.A303S and PYGL. In this study, mutation sites are denoted as “site in Protein” (e.g., A303S in PYGL), while mutant proteins follow HGVS standards (e.g., PYGL p.A303S); **g**, Effects of wild-type (WT) and mutant (MUT) PYGL overexpression on lung cancer cell growth. Top: Immunoblot validation of stable ectopic expression of WT and MUT PYGL in H1975 and PC-9 cells; GAPDH served as the loading control. Bottom: Proliferation curves of indicated cell lines monitored over time by CCK-8 assay. Values represent the mean ± SD (n=5). P values were calculated using two-way ANOVA; exact P values are displayed in the figure.

We next sought to identify mutations potentially associated with tumor development or progression. To this end, we evaluated the candidates from two complementary perspectives: their abundance and clinical relevance in the LUAD cohort, and their predicted effects on protein stability and function. First, using the site-resolved mutation abundance matrix, we examined differential abundance between paired tumor and adjacent tissues, associations with patient survival, and recurrence across individuals (**Fig. 4c**). Notably, approximately 1.0% (n = 53) of the identified mutations were significantly correlated with unfavorable clinical outcomes (overall survival (OS) or disease-free survival (DFS)) in patients. And the proportion of mutations detected in at least 20 patients was 1.8% (n = 93) Second, we performed mutation-effect analysis to assess the potential structural and functional consequences of the candidate amino-acid substitutions (**Fig. 4d**). Mutated protein stability predicted by I-Mutant2.0^51^ indicated that 86.9% (n = 4,433) of the mutations would decrease protein thermodynamic stability, while only 13.1% (n = 670) were predicted to stabilize the protein structure. AlphaMissense^52^ and PolyPhen-2^53^ represent the application of deep learning models and traditional statistical classifiers, respectively, in mutation effect prediction.

By integrating abundance, clinical recurrence, survival association, predicted thermodynamic stability and predicted functional impact, we prioritized 76 unique mutations with potential relevance to tumor development or progression (**Fig. 4e**). These candidates were distributed across four partially overlapping evidence categories: (1) destabilizing deleterious differentially expressed mutations (n = 48, red); (2) stabilizing deleterious differentially expressed mutations (n = 4, yellow); (3) destabilizing deleterious mutations associated with survival (n = 16, blue); and (4) high-frequency destabilizing deleterious mutations (occurrence ≥ 20, n = 22, purple). The overlap among these categories reflects mutations supported by multiple independent lines of evidence.

Among these potentially tumor-associated mutations, we next sought candidates with prognostic and therapeutic relevance that could be subjected to experimental validation. Mutation-based biomarkers are attractive because they capture tumor-specific changes in protein function that cannot be inferred from total protein abundance alone, and recurrent mutations are more likely to reflect functionally selected, clinically relevant events. We further reasoned that a mutant form whose prognostic association diverges from, or opposes, that of its wild-type counterpart would be particularly informative: such a pattern argues against a simple abundance-driven effect and points to mutation-specific biological consequences. We reasoned that a mutant form displaying a tumor-associated pattern divergent from, or opposite to, that of its corresponding wild-type form might provide biological insights beyond changes in total protein abundance.

On this basis, we prioritized PYGL p.A303S and CALR p.P216L as candidates because their mutant and wild-type forms exhibited opposing associations with overall survival, and their mutation frequencies exceeded 30%. Specifically, high expression of wild-type PYGL correlated with poor OS in lung adenocarcinoma, whereas its mutation form was associated with favorable outcomes (**Fig. 4f**). Conversely, wild-type CALR served as a favorable prognostic indicator for OS, while its mutant counterpart predicted adverse clinical outcome. To functionally validate these prioritized candidates, we performed orthogonal validation using well-established LUAD cell lines, H1975 and PC-9. Consistent with a previous report^54^, stable overexpression of wild-type PYGL markedly accelerated cell proliferation, whereas ectopic expression of the PYGL p.A303S mutant significantly suppressed tumor cell growth (**Fig. 4g**). Collectively, these findings highlight PYGL p.A303S as a promising prognostic biomarker candidate and functional modulator in lung adenocarcinoma.

We additionally investigated whether local structural and energetic perturbations could provide an orthogonal basis for prioritizing cancer-relevant mutations. Specifically, for each mutation, FoldX^55^ was used to generate a single-point mutant from the corresponding wild-type structure, and PyMOL^56^ was to quantify folding free-energy change (ΔΔG), relative solvent accessibility (RSA) and solvent-accessible surface area change (ΔSASA) at the mutated site, all-atom RMSD within a 6 Å neighborhood of the mutated residue, changes in residue contact number within 4.5 Å (Δcontact), and changes in hydrogen-bond count within 3.5 Å (Δhydrogen bond). Across the four high-priority mutation categories, structural and energetic perturbations were concordant: mutations with increased local RMSD often exhibited elevated ΔΔG, reduced site-level SASA, and remodeled contact networks, suggesting coupled conformational and stability changes in functional microenvironments. Higher AlphaMissense or PolyPhen-2 scores were likewise associated with greater energetic and local structural perturbation. We further related eight prioritized mutations to malignancy through the wild-type functions of their corresponding proteins. These mutations included HBA1 p.F47E, ANXA2P2 p.I129D, NUCB1 p.F138G, PSMA6 p.T73Y, SPPL2A p.V88M, ME3 p.Y200D, POTEE p.K913G, and ATP5F1A p.V443G. Notably, HBA1 p.F47E exhibited the largest local structural deviation among the identified substitutions (local RMSD = 1.08 Å) and a predicted increase in folding free energy (ΔΔG = +3.17 kcal/mol), suggesting reduced protein stability. HBA1 encodes the α-globin subunit of hemoglobin, which is essential for oxygen transport. F47 is located near the heme-binding pocket, where a previously reported F-to-S substitution has been associated with hemoglobin instability^57^. Therefore, F47E may disrupt the local heme environment and compromise hemoglobin function, potentially affecting oxygen-related processes and redox homeostasis in LUAD. Furthermore, NUCB1 p.F138G showed a pronounced predicted destabilizing effect (ΔΔG = +4.49 kcal/mol), accompanied by a modest decrease in solvent-accessible surface area (ΔSASA = −3.18 Å²) and local structural deviation (local RMSD = 0.50 Å). NUCB1 is a Golgi-associated calcium-binding protein involved in calcium-dependent trafficking of matrix metalloproteinases, including MMP2 and MT1-MMP, which are important regulators of extracellular matrix remodeling and cell invasion^58^. The substitution of F138G may weaken local structural packing and compromise protein stability, potentially affecting NUCB1-mediated protein trafficking and invasion-related processes in LUAD.

Together, these analyses show that π-SPECFormer can convert previously unassigned clinical proteomic spectra into experimentally testable biological hypotheses. Starting from 5,103 candidate mutation peptides, the framework integrated spectral evidence, paired quantitative proteomics, population and disease annotation, clinical outcomes, mutation-effect predictions and local structural analyses to identify PYGL p.A303S as a previously unreported functional and prognostic candidate mutation in LUAD. In parallel, mutations prioritized on the basis of coupled energetic and local structural perturbations provide additional candidates for future mechanistic validation. More broadly, these findings show that proteomic dark matter contains clinically informative molecular variation that were missed by conventional reference-database-centered analyses.

### Mutation-remodeled protein interfaces reveal opportunities for targeted drug design

Having identified candidate mutations with tumor-associated abundance, clinical relevance and predicted structural effects, we next investigated whether these substitutions could alter interactions with established oncology drug targets. We reasoned that a mutation located within or adjacent to a protein–protein interface could remodel local geometry, electrostatics or solvent accessibility, thereby affecting oncogenic signaling or treatment response. Such mutation-remodeled interfaces might also create therapeutic opportunities: compounds designed to occupy the altered interface, stabilize or disrupt the affected complex, or selectively recognize a mutation-induced pocket could preferentially target the mutation protein while sparing its wild-type counterpart.

As shown in **Fig. 5a**, leveraging the transformative capability of AlphaFold 3 (AF3) in modeling protein assemblies^59, 60^, we established a structural interaction discovery platform to identify novel interfaces between the cancer-related mutations identified in this study and established oncology drug targets. This approach is predicated on the rationale that antineoplastic agents typically target proteins governing critical tumorigenic processes; consequently, mutations within these targets or their interacting partners can profoundly modulate therapeutic efficacy or disease progression. We curated a comprehensive library of drug targets by extracting 235 drugs from 246 FDA-approved cancer treatment regimens (1998–present, sourced from OncoKB^61^) and mapping them to 276 targets via DrugBank^62^. Ultimately, 248 human protein targets were selected for downstream complex prediction. We systematically evaluated the interactome between these 248 drug targets and 69 wild-type proteins spanning four mutation categories (76 unique mutations in total) (**Fig. 5a**).

**Fig. 5.**
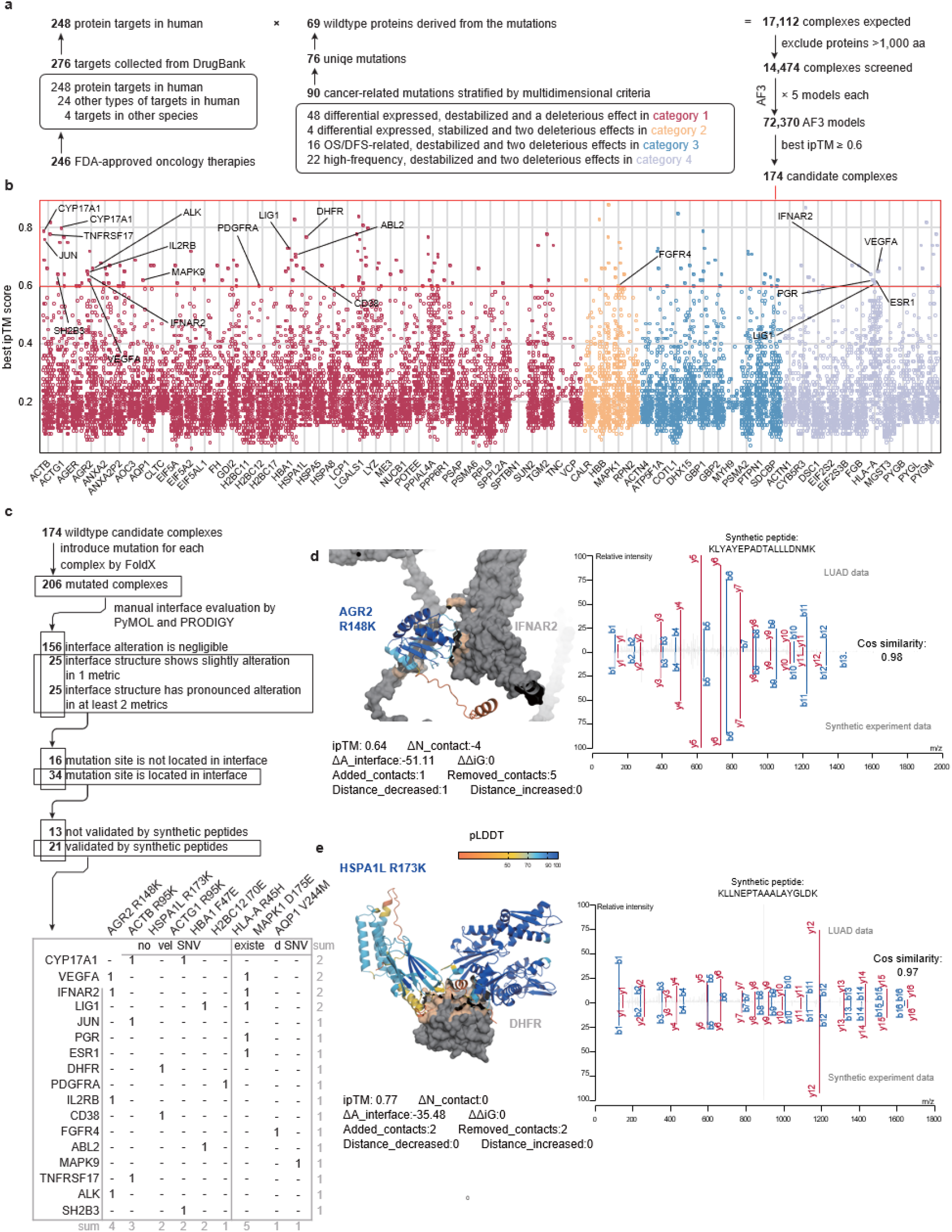
Systematic screening of mutation perturbations on the interaction landscape of FDA-approved anticancer drug targets. **a**, Schematic overview of the workflow designed to characterize interactions between mutated proteins and FDA-approved drug targets. **b,** Distribution of the highest ipTM scores for 14,474 complexes formed by the wild-type counterparts of 69 mutated proteins. Complexes are colour by mutation category, with 174 high-confidence candidates (ipTM ≥ 0.6) highlighted. **c,** The workflow for screening candidate mutant protein-drug target interaction models. Mutant models were generated using FoldX. Candidates were filtered based on structural metrics calculated via PyMOL and Prodigy—including changes in average contact number (ΔN_contact), interface surface area (ΔA_interface), and binding free energy (ΔΔiG)—combined with interface conformational analysis, synthetic peptide validation, and literature curation. **d and e**, Predicted structural and interface-contact changes in representative mutation protein–drug-target complexes involving AGR2 R148K (d) and HSPA1L R173K (e), together with comparisons between the corresponding cohort-derived and synthetic-peptide MS/MS spectra.

To reduce redundant computation, we implemented an optimized AF3 workflow in which multiple-sequence alignments were calculated once for each monomer and reused across complex predictions. Protein pairs containing a component longer than 1,000 amino acids were excluded, leaving 14,474 pairs for structural screening. Of these, 174 complexes achieved a best interface predicted template modeling score (ipTM) of at least 0.6 and were retained as candidate interactions (**Fig. 5b**). Candidate complexes were most frequently observed among proteins in the first mutation-prioritization category, although this enrichment was influenced by the number of proteins screened in each category and therefore does not alone indicate greater biological relevance. Subsequently, we modeled the specific mutations into these 174 complexes using FoldX, generating 206 mutant protein-drug target complex models (**Fig. 5c**).

To quantify the structural impact of these mutations, we developed a custom visualization and analysis pipeline. Complex structures were rendered with the drug target in gray and the interaction interface in light orange, while the mutant protein was colored by pLDDT scores. We categorized the 206 mutant complexes into three groups by comparing them to their wild-type counterparts based on the interface contact number (ΔN_contact), buried interface area (ΔA_interface), and binding affinity change (ΔΔ^i^G). Group 1 (n = 156) exhibited negligible structural perturbation (all z-score normalized metrics between −1 and 1). In contrast, Group 2 (n = 25) and 3 (n = 25) exhibited slight and pronounced alterations, respectively, with one metric and at least two metrics exceeding the threshold (**Fig. 5c**). We therefore focused our downstream analysis on the 50 complexes in Groups 2 and 3 that exhibited significant interface remodeling. We excluded 16 models where the mutation site was located outside the binding interface, as these are less likely to directly disrupt interaction mechanics. The remaining 34 complexes, featuring mutations directly at the binding interface, were prioritized for validation. To ensure the accuracy of the mutation sites, we conducted synthetic peptide experiments and compared the similarity between the synthetic peptide experiment spectra and the original spectra. We identified nine mutations (corresponding to 21 complexes), and the associated mutant peptides were validated by their high similarity to experimentally generated synthetic-peptide spectra. Consequently, we highlighted these 9 mutations to be high-confidence mutations with candidate tumor relevance and their dynamic changes in interface contacts. We found that 6 remained unreported after cross-referencing dbSNP, GDC, COSMIC, and PubMed literature, whereas 3 were independently corroborated by these external reference resources. These findings suggest that our model can both uncover novel candidate disease-relevant SNVs and reliably recover previously documented mutations, providing external support for its accuracy.

To further elucidate the potential impact of these mutations on protein–drug target interactions, we quantified the dynamic changes in interface contacts at the binding interfaces of candidate complexes (**Fig. 5d-e**). Specifically, by comparing each mutant protein–drug target complex with its corresponding wild-type one, interface contacts were categorized into four classes: (1) added contacts; (2) removed contacts; (3) shared contacts with decreased contact distances; and (4) shared contacts with increased contact distances. The AGR2 is a disulfide isomerase playing a role in cell migration, differentiation and growth. AGR2 p.R148K represents a conservative amino acid substitution within a ligand-responsive region (residues 138–149) adjacent to the peptide-binding loop of AGR2. Previous studies have demonstrated that this region undergoes conformational changes upon ligand binding, while AGR2 interacts with EpCAM and facilitates EGFR trafficking to the cell surface^63^. R148K potentially affecting AGR2-mediated protein recognition in LUAD. AGR2 p.R148K mutation affected large number of interaction models (n = 4), and the predicted AGR2 p.R148K–IFNAR2 complex exhibited significant changes in interface contact (**Fig. 5d**). HSPA1L, a member of the Hsp70 (70 kDa heat shock protein) family, showed a limited number of interaction models affected by the R173K mutation (n = 2) in the N-terminal nucleotide-binding domain (NBD), accompanied by moderately evident changes in interface contacts. Nevertheless, prior studies have reported that Hsp70 family chaperones can recognize mutation-exposed DHFR and facilitate its clearance via the proteasome pathway^64^. Therefore, the R173K substitution may alter local interactions within the NBD and potentially affect chaperone regulation, with possible implications for protein homeostasis and cancer-associated signaling in LUAD. (**Fig. 5e**).

We assembled the spectrally supported substitutions, candidate interaction partners, AF3 complex models, interface-confidence scores, predicted energetic changes and residue-level contact annotations into a mutation–drug-target interface resource. This resource is intended to support hypothesis generation rather than to provide a catalogue of experimentally confirmed interactions. It enables researchers to prioritize candidate complexes according to spectral confidence, clinical association, interface localization and predicted structural perturbation, and to retrieve paired wild-type and mutation models for further analysis.

Importantly, the mutation-centered regions represented in this work may provide structural starting points for future targeted drug discovery. For substitutions predicted to strengthen an oncogenic interaction, the surrounding interface could be explored for molecules that selectively disrupt the mutant complex. Conversely, substitutions predicted to weaken a potentially protective interaction could motivate the design of interface stabilizers. Depending on interface topology and cellular localization, subsequent studies could evaluate small molecules, fragments, constrained peptides, macrocycles or biologics directed towards the mutation-containing region. The paired wild-type and mutation models may also help prioritize designs that recognize structural features specific to the mutation while minimizing perturbation of the corresponding wild-type protein.

These potential applications remain prospective. AF3 confidence scores, FoldX energetic estimates and contact changes do not demonstrate that the predicted complexes form in cells, that the substitutions alter binding affinity or that the mutation-centered regions are pharmacologically tractable. Future work will require confirmation by co-localization or co-immunoprecipitation, quantitative binding assays, mutational scanning, molecular-dynamics simulations and experimental assessment of pocket or interface accessibility before compound discovery is undertaken.

Collectively, these analyses establish a spectra-supported structural resource connecting candidate proteomic mutations with predicted interfaces involving established oncology drug targets. Although the interactions and their functional consequences remain to be experimentally validated, the resource provides residue-level hypotheses and paired wild-type and mutation structural models for community use. In particular, mutation-centered interface regions highlighted by the analysis offer starting points for future structure-guided development of small molecules, peptides or biologics designed to selectively modulate mutation-associated protein interactions.

## Discussions

In conventional mass-spectrometry-based proteomics, nearly 50% of high-quality spectra remain uninterpreted. These unannotated spectra carry substantial biological significance, encoding information on single amino acid polymorphisms, alternative splicing events, diverse PTMs and the translation products of non-canonical open reading frames — collectively constituting a vast and largely uncharted proteomic dark matter.

In recent years, deep learning has been introduced into spectral data analysis, emerging as a transformative technical force in this field. Nevertheless, most existing task-specific models rely predominantly on supervised training with peptide labels generated from database-search-derived PSMs — giving rise to an intractable computational paradox: the algorithm aspires to discover unknown biological variation, yet its epistemic horizon is strictly bounded by the biases inherent in existing reference annotations. To address these challenges, we present π-SPECFormer, a novel computational paradigm and foundational model purpose-built for mass spectrometry data. During pre-training, the model autonomously learns — via a contrastive learning objective — to capture the physicochemical fragmentation patterns of amino acid sequences, fragment ion intensity distributions, and high-dimensional spectral patterns directly from hundreds of millions of MS/MS spectra. This unbiased representation learning capability enables π-SPECFormer to transcend the boundaries of canonical proteomics, acquiring a genuine understanding of the “underlying language” of dark matter. Application of π-SPECFormer to mutation discovery further expanded our view of the molecular heterogeneity of lung cancer and, more importantly, provided evidence that proteomic dark matter can contain biologically relevant molecular variation.

π-SPECFormer has limitations. The first lies in the limited diversity of training data. Expanding to a broader variety of data, such as more species or data-independent acquisition (DIA) data, to enhance the model’s generalization is a future research direction. As we include more data, balancing the model’s ability to differentiate distinct data features while identifying similarities between datasets is a crucial area for further investigation.

The second is the experimental validation. In this study, we identified a substantial number of novel mutations and unannotated peptides in the LUAD cohort that are potentially associated with tumor initiation and progression. However, owing to constraints in time and resources, validation of these dark matter candidates currently relies primarily on computational false discovery rate control and structural simulation. Directions of experimental validation are warranted in future work: (a) experimental validation of the binding relationships between mutant and wild-type proteins — for instance, through co-immunoprecipitation (Co-IP) assays — to address potential false positives arising from the AlphaFold3-based complex confidence scoring step; and (b) functional characterization of mutation proteins in tumor-relevant biological processes, including proliferation and metastasis, through in vivo cellular models.

Investigating the biological significance of these unidentified spectra holds substantial potential. Dark matter may represent unknown or newly discovered proteins, translation products from non-coding regions, mutation proteins within specific organisms, novel splice mutations, or unexpected post translationally modified peptides. Additionally, the dark matter may also contain those low-abundance proteins that could play key roles in physiological or pathological processes. Moreover, analyzing dark matter could aid in identifying novel disease biomarkers or drug targets, offering new insights into disease mechanisms and advancing the development of precision medicine. We are confident that π-SPECFormer and its future versions will become a powerful tool for proteomic data analysis, especially for dark proteome mining.

## Methods

### Pre-training and down-stream tasks

#### Pre-training Dataset and pre-processing

In this study, to construct large-scale datasets for model pre-training, we built two primary datasets. The first dataset consists of unidentified spectra (whether they can be identified using traditional database search methods), containing 178,171,669 spectra after preprocessing. The second dataset consists of identified spectra, or PSM, which have been successfully matched to corresponding peptide sequences through database searches. This dataset contains 113,737,306 spectra after preprocessing. The spectra were grouped according to peptides and their charge states, and groups with only one spectrum were removed, resulting in the creation of 2,315,858 spectrum groups, with an average of 49 spectra per group. For the two spectral datasets, we applied the following preprocessing steps: (1) Selection of spectra with precursor ion charges between +2 and +6: This charge range represents the most commonly observed precursor ion charge states in mass spectrometry. (2) Selection of spectra with 20 to 500 signal peaks: This criterion was set because spectra with fewer than 20 signal peaks are likely the result of incomplete peptide fragmentation, leading to insufficient meaningful peaks. On the other hand, spectra with more than 500 signal peaks may contain excessive noise due to overly extensive peptide fragmentation.

#### The peak encoder of π-SPECFormer

As shown in the following formula, to encode the spectral data, we employed a method similar to Transformer positional encoding for encoding the spectral data.

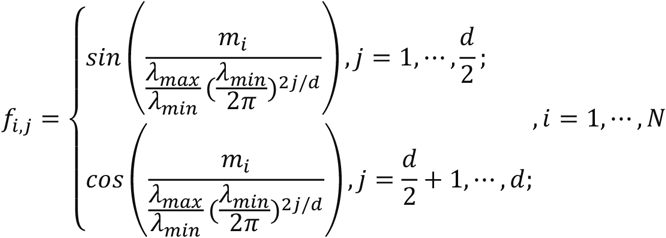

In this context, *λ_min_* = 0.001, *λ_max_* = 10,000 and *m_i_* represents the mass-to-charge ratio (m/z) of the i-th signal peak in the spectrum, while *f_i_*_,*j*_ represents the j-th dimension of the i-th signal peak in the spectrum. The variable *d* refers to the dimensionality mapped by the model. Additionally, for the intensity information in the spectral data, since its precision is lower than that of the mass-to-charge ratio, we map it to dd dimensions using a linear layer. This mapped representation is then added to the representation derived from the mass-to-charge ratio and subsequently fed into the model.

#### The Momentum Encoder

In this model, there are two Spectrum encoders: one for extracting features from the query spectrum (the online encoder) and the other for extracting features from spectra in the queue (the momentum encoder). During each training iteration, the parameters of the momentum encoder are updated in proportion (usually by a small factor) to the parameters of the online encoder. The update of the momentum encoder follows the rule shown in equation (1).

#### Pre-training strategy

In Stage 1, we believed the model has sufficiently learned the internal distribution patterns of fragment ions within spectra. Inspired by the development path of large language models—where early models like GPT-1 through GPT-3 relied on self-supervised pre-training on text data (similar to our Stage 1)—GPT-4 built on this foundation by directly utilizing task-specific conversational data, achieving substantial performance gains. Given that our downstream task aims to elucidate the correspondence between spectra and peptide sequences, we directly leveraged PSMs (Peptide-Spectrum Matches) data for further pre-training to enhance the model. Notably, while peptide sequence information is incorporated only during data preprocessing, the actual model training remained solely based on spectral data.

#### Pre-training parameters

In terms of model construction, this model consists of 9 Transformer modules with an embedding dimension of 512 and 8 attention heads. Under this framework, the model contains a total of 21.07 million learnable parameters.

For model training: (1) π-SPECFormer-stage1 pre-training: During pre-training with unlabeled spectra, the batch size was set to 256, the learning rate was 0.03, and the number of training epochs was 10. The masking ratio for spectrum signal peaks was 30%. The model was trained for a total of 13 days and 1 hour using 4 V100 Tensor Core GPUs. (2) π-SPECFormer-stage2 pre-training: For further pre-training with labeled spectra, we used the same hyperparameter settings as above. This stage of the model was trained for a total of 3 days and 18 hours using 32 V100 Tensor Core GPUs.

#### Evaluation metrics

π-SPECFormer employs the same evaluation metrics as other *de novo* sequencing models: amino acid precision (AA precision), amino acid recall (AA recall), and peptide recall.

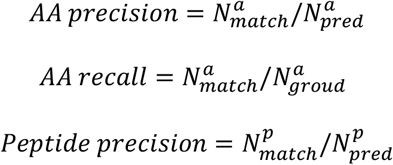

In this context, *N^a^_match_*, refers to the total number of amino acids predicted by the model that match the ground truth labels, *N^a^_pred_* represents the total number of amino acids predicted by the model, and *N^a^_groud_* is the total number of amino acids in the ground truth labels.

Similarly, *N^p^_match_* denotes the total number of peptide sequences predicted by the model that exactly match the ground truth sequences, where even a single mismatch in any position renders the prediction incorrect. *N^p^_pred_* represents the total number of peptide sequences in the ground truth labels.

#### Mutation mining in cancer cohort data

##### Pipeline for mining mutational dark matter

To mine mutational dark matter, we implemented a five-stage workflow to identify previously unannotated mutations. First, to better match the characteristics of our dataset, the π-SPECFormer model was fine-tuned using high-confidence peptide–spectrum matches (PSMs) derived from the LUAD cohort. Next, de novo sequencing was performed on spectra that remained unidentified by conventional database searching (“dark matter”), followed by stringent quality control to retain only sequences with a precursor mass error within 100 ppm and a prediction confidence score >70. The resulting de novo sequences were then aligned against the human reference proteome, and peptides harboring exactly one amino acid substitution were retained as putative mutant peptides. Subsequently, the abundances of these mutant peptides were quantified using the PANDA algorithm to generate a mutation quantification matrix. Finally, this matrix was used for systematic statistical and functional downstream analyses to characterize the biological significance of the identified mutations.

##### Classification of Mutation by Genetic Polymorphism

To characterize the genetic polymorphisms of the 5,103 mutations, we performed a comprehensive analysis of mutation frequency and database coverage. Each mutation was cross-referenced against major public repositories, including dbSNP (https://www.ncbi.nlm.nih.gov/snp/), GDC (https://portal.gdc.cancer.gov/), COSMIC (https://cancer.sanger.ac.uk/cosmic), and literature indexed in PubMed (https://pubmed.ncbi.nlm.nih.gov/). Mutations were categorized as common single nucleotide polymorphisms (SNPs), known single nucleotide mutations (SNVs), or novel SNVs based on their minor allele frequency (MAF) and documentation status. Specifically, mutations recorded in dbSNP with a MAF > 1% in at least two genomic sequencing datasets were classified as common SNPs, following the guidelines of the 1000 Genomes Project^53^.

##### Prediction of Mutation Effects

We assessed the structural and functional impact of the identified mutations using a multi-tool framework. Thermodynamic stability changes upon mutation were predicted with I-Mutant2.0 in Sequence mode using the corresponding amino acid sequences with default parameters. Functional impact was evaluated using AlphaMissense scores retrieved from the pre-computed AlphaMissense dataset (https://zenodo.org/records/8360242) and PolyPhen-2 predictions generated via the PolyPhen-2 web server using the batch query function with the HumVar model under default settings. The resulting stability and pathogenicity annotations were integrated for downstream prioritization and stratification of mutations based on predicted effect, genetic properties, and clinical association.

##### Structural Modeling of Mutant Proteins

For each mutation, the corresponding wild-type experimental structure was preferentially selected from PDBe best_structures resource (https://www.ebi.ac.uk/pdbe/api/mappings/best_structures/). Candidate entries were first filtered to retain only those with UniProt residue-level coverage of the mutated position. The remaining structures were then ranked by experimental method (X-ray diffraction > neutron diffraction > electron microscopy/cryo-EM > others) and, within the same method category, by increasing resolution (lower Å values preferred); ties were further resolved by higher sequence coverage. When no experimental structure covered the mutation site, the AlphaFold Protein Structure Database model (v6; https://alphafold.com/) was used as a substitute. Structural refinement and in silico mutagenesis were performed in FoldX v5.1 using default parameters: all starting structures were first optimized with RepairPDB, followed by introduction of the desired substitutions using BuildModel. Each mutation was modeled in five independent runs for monomeric proteins and in three independent runs for AF3-predicted protein–drug target complexes due to computational cost, and interface-related metrics were averaged across repeats. All wild-type and mutant models were visualized and inspected in PyMOL.

##### Evaluation of local structural perturbations

To quantify mutation-induced changes in local structure and thermodynamics for monomeric (non-complex) models, we compared each mutant with its wild-type counterpart using several residue-centric metrics computed in PyMOL (v3.2.0a, open-source) under fixed geometric rules: (1) folding free-energy change (ΔΔG), taken from the FoldX output for each model; (2) relative solvent accessibility (RSA) at the mutated residue, obtained with get_sasa_relative; (3) change in solvent-accessible surface area (ΔSASA) at the mutated residue, computed as the difference in residue-level SASA between mutant and wild-type using get_area (with identical probe settings across pairs); (4) local all-atom RMSD within a 6 Å neighbourhood of the mutated residue after superposing matched heavy atoms in that neighbourhood; (5) change in residue contact number (Δcontact), defined as the difference in the number of intra-chain partner residues with any heavy-atom contact to the mutated residue within a 4.5 Å cutoff (excluding self-terms); and (6) change in hydrogen-bond count (Δhydrogen bond), defined as the difference in the number of putative hydrogen-bonding interactions between atoms of the mutated residue (subject to a donor/acceptor atom-type screen) and heavy atoms on other same-chain, non-neighbour residues within 3.5 Å—using a 3.5 Å cutoff for most pairs and a 4.0 Å cutoff when a sulfur atom was involved on either side. Because these descriptors have different scales, we defined high local-perturbation events within the analyzed monomer cohort by applying, to each metric separately, a magnitude threshold equal to the 75th percentile of its absolute values (i.e., approximately the top quartile by magnitude), enabling cross-metric comparison without treating these cutoffs as universal significance tests. In addition, we juxtaposed the same mutations with sequence/function-level deleteriousness predictors (e.g., AlphaMissense and PolyPhen-2); given multi-step mutation enrichment and limited sample size, such comparisons are exploratory and not presented as primary statistical endpoints. All extractions were implemented in batch with fixed scripts and recorded cutoffs to ensure reproducibility. Curation of FDA-Approved Drug Targets

We curated a set of clinically relevant drug targets by screening FDA-approved oncology therapies from OncoKB (v5.1; https://www.oncokb.org/), which comprised 246 FDA-approved cancer treatment regimens (including six subsequently withdrawn therapies) spanning 1998 to the present. Combination regimens were decomposed into individual drugs, yielding 235 unique antineoplastic drugs. Drug-target annotations were retrieved from the DrugBank database (v5.1.12; https://go.drugbank.com/) by extracting entries labeled as targets, resulting in 276 unique targets. After restricting to human proteins, 248 unique protein targets were retained for downstream complex prediction. Amino acid sequences were obtained from UniProt (https://www.uniprot.org/) through ID mapping.

##### AlphaFold 3-based Protein Complex Prediction

Protein complex structures were modeled using AlphaFold 3 (AF3; v3.0.1). To improve computational efficiency, we decoupled database search/MSA generation from complex structure prediction: for each monomeric chain (i.e., the cancer-associated mutant protein and the drug target), the AF3 Search Only stage was first executed to generate multiple sequence alignments (MSAs) and template information using JackHMMer v3.3 against the Big Fantastic Database (BFD), MGnify protein clusters, the RNA subset of NCBI nt, PDB seqres, Rfam, the active-sequence subset of RNAcentral, UniProtKB, and UniRef90. The resulting per-chain MSA/template outputs were then reused to construct a complex-level JSON input, in which chain-specific evolutionary information was combined to enable subsequent AF3 complex prediction; this assembled input was passed to the AF3 Predict Only module to obtain the 3D structure of each complex. From the standard AF3 outputs, we extracted per-residue confidence scores (pLDDT), predicted aligned error (PAE) matrices, predicted TM-score (pTM), predicted interface TM-score (ipTM), and the model ranking score; for each complex, the top-ranked model (highest ranking score) was selected for downstream analyses. When the same monomer appeared across multiple predicted complexes, its confidence metrics were summarized by averaging residue-level or complex-level scores across all relevant predictions. All amino acid sequences for proteins were derived from the Homo sapiens reference proteome (Proteome ID: UP000005640).

##### Evaluation of Binding Interface Alterations

To quantify the structural impact of mutations on protein–protein interactions, we compared each mutant complex with its corresponding wild-type counterpart using three interface-level metrics: (1) change in interface contact number (ΔN_contact), defined as the difference in the number of inter-chain atomic contacts within a 4.5 Å cutoff at the binding interface and computed using PyMOL (v3.2.0a, open-source); (2) change in buried interface area (ΔA_interface), defined as the difference in solvent-accessible surface area (SASA) buried upon complex formation and computed in PyMOL; and (3) change in binding affinity (ΔΔiG), defined as the difference in predicted binding free energy estimated with PRODIGY (v2.4.0) using default parameters. For visualization, complex structures were rendered with the drug target in gray and the interaction interface in light orange, while the mutant protein was colored by pLDDT scores.

### Synthetic Peptide Experiment

#### LC-MS/MS Analysis of Synthetic Peptide Experiment

Chromatographic separation was performed on an EASY-nLC 1200 system (Thermo Fisher Scientific). A total of 50 ng (5 μL) of peptides was loaded onto a custom-packed trap column (2 cm × 100 μm i.d.) packed with 1.9 μm C18-AQ resin (Dr. Maisch GmbH). Peptides were separated on a custom-packed analytical column (30 cm × 150 μm i.d.) packed with ReproSil-Pur C18-AQ resin (1.9 μm; Dr. Maisch GmbH). Mobile phase A consisted of 0.1% formic acid in water, and mobile phase B consisted of 0.1% formic acid in 80% acetonitrile. The flow rate was maintained at 600 nL/min. A 30-min gradient was applied as follows: 7–12% B (0–2 min); 12–32% B (2–13 min); 32–45% B (13–19 min); 45–95% B (19–21 min); and held at 95% B (21–30 min).

Mass spectrometry data were acquired using an Orbitrap Fusion Lumos Tribrid mass spectrometer (Thermo Fisher Scientific) operating in positive ion mode. The ion transfer tube temperature was set to 320 °C, and the spray voltage was 2.2 kV. The instrument was operated in OT-OT mode. MS1 full scans were acquired in the Orbitrap across a mass range of 350–1500 m/z at a resolution of 120,000 (at 200 m/z), with an automatic gain control (AGC) target of 1 × 10⁶ and a maximum injection time of 50 ms. For MS2, the instrument operated in Top-Speed mode with a cycle time of 3 s. Precursor ions with charge states of 2–7 were selected for fragmentation via High-energy Collisional Dissociation (HCD) at a normalized collision energy (NCE) of 32%. MS2 spectra were acquired in the Orbitrap at a resolution of 15,000, with a fixed first mass of 120 m/z. The AGC target was set to 5 × 10⁴ with a maximum injection time of 22 ms. A dynamic exclusion window of 25 s was applied to minimize repeated sequencing of abundant precursors.

#### Database Searching for Synthetic Peptide Experiment Data

Raw mass spectrometry data were processed and searched using Thermo Proteome Discoverer (version 2.4.1.15). Spectra were queried against a composite database containing the human proteome (20,420 entries) and 32 synthetic peptide sequences. Trypsin (Full) was specified as the proteolytic enzyme, allowing for a maximum of two missed cleavage sites. The peptide length range was restricted to 6–144 amino acids. Mass tolerances were set to 15 ppm for precursor ions and 0.05 Da for fragment ions. Carbamidomethylation of cysteine (+57.021 Da) was defined as a static modification. Dynamic (variable) modifications included oxidation of methionine (+15.995 Da), phosphorylation of serine, threonine, and tyrosine (+79.966 Da), and N-terminal acetylation (+42.011 Da).

### Validation experiment for candidate mutations

#### Cell culture

The human embryonic kidney cell line HEK293T (RRID:CVCL_0063) and the human lung adenocarcinoma cell lines PC-9 (RRID:CVCL_B260) and NCI-H1975 (RRID:CVCL_1511) were obtained from our laboratory cell bank. HEK293T and PC-9 cells were cultured in high-glucose Dulbecco’s modified Eagle’s medium (DMEM; Gibco), whereas NCI-H1975 cells were maintained in RPMI-1640 medium (Gibco). All culture media were supplemented with 10% fetal bovine serum (FBS; NEWZERUM, #FBS-UE500) and 1% penicillin–streptomycin (100 U/mL penicillin and 100 μg/mL streptomycin; Gibco, #15140-122). Cells were incubated at 37 °C in a humidified atmosphere containing 5% CO₂. All cell lines were recently authenticated by short tandem repeat (STR) profiling and routinely tested to confirm the absence of mycoplasma contamination.

#### Generation of stable cell lines

Coding sequences for human wild-type (WT) PYGL (NCBI reference sequence: NM_002863.5) and CALR (NM_004343.4), as well as their respective mutant counterparts (PYGL p.A303S and CALR p.P216L), were cloned into the pHAGE lentiviral expression vector (Addgene, #118692) with a C-terminal Flag tag. For lentivirus packaging, HEK293T cells were co-transfected with the corresponding pHAGE transfer plasmid alongside the packaging plasmids psPAX2 (Addgene, #12260) and pMD2.G (Addgene, #12259) using polyethyleneimine (PEI; Polysciences, #24765). Lentiviral supernatants were harvested 48 h post-transfection, passed through a 0.45-μm filter, and subsequently used to transduce H1975 and PC-9 cells in the presence of 8 μg/mL polybrene (Solarbio, #H8761). At 24 h post-infection, cells were subjected to selection with 5 μg/mL puromycin (Solarbio, #P8230) for two weeks to establish stable polyclonal populations. Ectopic expression was confirmed by Western blot analysis.

#### Western blot analysis

Indicated cells were lysed in RIPA buffer supplemented with a protease inhibitor cocktail (MCE, #HY-K0010). Cell lysates were sonicated and centrifuged at 12,000 × *g* at 4 °C for 15 min to remove insoluble debris. Equal amounts of protein were mixed with SDS loading buffer and denatured at 95 °C for 5 min. Protein samples were resolved by 10% SDS–PAGE and transferred onto 0.45-μm nitrocellulose membranes (Cytiva, #10600079). After blocking with 5% non-fat milk in TBST, membranes were incubated with HRP-conjugated primary antibodies against Flag (Sigma-Aldrich, #A8592; 1:2,000) and GAPDH (Proteintech, #HRP-60004; 1:5,000). Immunoreactive bands were visualized using an enhanced chemiluminescence (ECL) substrate (NCM Biotech, #P10300) on an Amersham Imager 800 (Cytiva). Representative blots from three independent biological replicates are shown.

#### Cell proliferation assay

Cell proliferation was evaluated using the Cell Counting Kit-8 (CCK-8; Analysis Quiz, #AQ308) according to the manufacturer’s instructions. Briefly, stably transduced H1975 and PC-9 cells expressing wild-type or the indicated mutant constructs were seeded into 96-well plates at densities of 3,000 and 5,000 cells per well, respectively. Cell viability was monitored at 24-h intervals (0, 24, 48, and 72 h). At each designated time point, 10 μL of CCK-8 reagent (10% v/v) was added to each well, followed by incubation at 37 °C for 2 h. Absorbance at 450 nm was measured using a microplate reader (BioTek), and background absorbance from blank wells containing medium alone was subtracted. Statistical comparisons across groups and time points were performed using two-way ANOVA, with exact P values indicated directly in the corresponding panel.

## Author Contributions

Authors sharing identical initials are identified by their full names in parentheses. C.C. and Y.W. co-supervised and conceived the study. T.L. (Tianze Ling), F.X., X.H. and T.L (Tao Liu) collected preprocessed all the datasets for pre-training task, down-stream tasks and biological application. F.X. and T.L. (Tianze Ling) proposed the unsupervised pre-training model and implemented related pre-training tasks. T.L. (Tianze Ling), J.X., T.Y., Z.L. and T.L. (Tao Liu) implemented experiments of *de novo s*equencing task. T.L. (Tianze Ling) and T.L. (Tao Liu) implemented the *de novo* model training for biological applications. S.C., T.L. (Tianze Ling) and T.L. (Tao Liu) implemented the experiments of spectrum clustering task. T.L. (Tianze Ling), F.X. and T.L. (Tao Liu) performed experiments of PTMs detection task. T.L. (Tianze Ling), T.L. (Tao Liu) and Y.Z. constructed the pipeline of cancer-related mutations mining and performed the analysis. T.L. (Tao Liu) conducted the analysis of mutation effects and the modeling of mutant-drug target protein interactions. Y.Z. conducted the cell-based validation experiments for the mutations. R.X. provided guidance on the mutation mining. K.L. and G.W. performed the LC–MS/MS experiment analysis of the synthetic peptides. L.X., J.Z. and Z.L. provided guidance on algorithm designs. S.Z.L. and F.H. coordinated the study and provided critical suggestions. T.L. (Tianze Ling) wrote the initial manuscript, and T.L. (Tao Liu), F.X., J.X., S.C., L.X., R.X., Y.W. and C.C. revised it. All authors read and approved the final manuscript.

## Acknowledgement

This work was financially supported by the National Key Research and Development Program of China (2025YFA1309400 to L.X. and C.C.), the National Natural Science Foundation of China (32688201 to F.H., 62606430 and 623B2086 to J.X.), the CAMS Innovation Fund for Medical Sciences (CIFMS, 2019-I2M-5-063 to F.H. and C.C.), the Research Project of the State Key Laboratory of Medical Proteomics (SKLP-Y202401 to C.C.), CCF-GHFund (2026005 to J.X.), National Science and Technology Innovation - Major Program (2022ZD0115101 to S.Z.L). We appreciate Xian Liu and Ming He for their guidance and computational support in establishing the AlphaFold 3-based protein–protein interaction prediction platform.

